# Machine learning as a tool for predicting insincere effort in power grips

**DOI:** 10.1101/068494

**Authors:** Peter Hahn, Eren Cenik, Karl-Josef Prommersberger, Marion Mühldorfer-Fodor

## Abstract

**Background:** It was not possible to detect the common problem of insincere grip effort in grip strength evaluation until now. The usually used JAMAR dynamometer has low sensitivity and specificity in distinguishing between maximal and submaximal effort. The manugraphy system may give additional information to the dynamometer measurements used to assess grip force, as it also measures the load distribution of the hand while it grips a cylinder. Until now, the data of load distribution evaluation were analyzed by comparing discrete variables (e.g., load values of a defined area). From another point of view, the results of manugraphy measurements form a pattern. Analyzing patterns is a typical domain of machine learning.

**Methods:** We used data from several studies that assessed load distribution with maximal and submaximal effort. They consisted of 2016 total observations, including 324 patterns of submaximal effort. The rest were from grips with maximal effort. After preparation and feature selection, XGBoost machine learning was used for classification of the patterns.

**Findings:** After applying machine learning to the given data, we were able to predict submaximal grip effort based on the inherent pattern with a sensitivity of 94% and a specificity of 100%.

**Interpretation:** Using techniques from applied predictive modeling, submaximal effort in grip strength testing could be detected with high accuracy through load distribution analysis. Machine learning is a suitable method for recognizing altered grip patterns.

## Introduction

Evaluation of grip strength is a common measurement in orthopedic and hand surgery for assessing hand function disability (Mafi 2012). Besides, grip strength is a widely accepted parameter for general health issues (Chung et al. 2014). For various reasons, such as receiving financial benefits, avoiding return to work, or secondary gain from illness, patients may feign the loss of grip force. Exact numbers are not available, but experienced investigators estimate malingering or symptom exaggeration in up to 30% of cases (Mittenberg et al. 2002). The JAMAR dynamometer is the most frequently used tool for investigating grip force in clinical practice. There are several techniques for figuring out submaximal effort with the JAMAR, like the five-rung test, the rapid exchange grip test and the coefficient of variation. All of these tests have low sensitivity and low specificity (Sindhu et al. 2012). Other tests require specialized equipment or calculations (Shechtman, Hope, et al. 2007; Sindhu & Shechtman 2011). They may give better results, but either their sensitivity or specificity is low in the end. Combining the test with other measurements, such as muscle mass circumference and callusing of the hands, is highly dependent on subjective perception. An objective method with high sensitivity and high specificity would be of great value.

Measuring grip strength and grip pattern with a sensor system (manugraphy) provides a variety of data. In contrast to the JAMAR dynamometer, manugraphy measures the load applied by the whole contact area of the hand using a sensor matrix, and it provides a high-resolution grip pattern for each measurement (Fig.1). A previous study, which tested 54 healthy volunteers, compared both hands in terms of the load applied to each of seven anatomical areas using the Wilcoxon-test. The corresponding areas of both hands were the same or very similar if both hands performed maximum grip force. If one hand was used with submaximal force, there was a statistical significant difference in 5/7 areas (left hand submaximal) or 7/7 areas (right hand submaximal), respectively (unpublished). After analyzing the data through statistical methods, the study showed the difference between maximal and submaximal effort for the whole study group, though it was impossible to predict insincere effort for a single patient.

**Fig 1:**
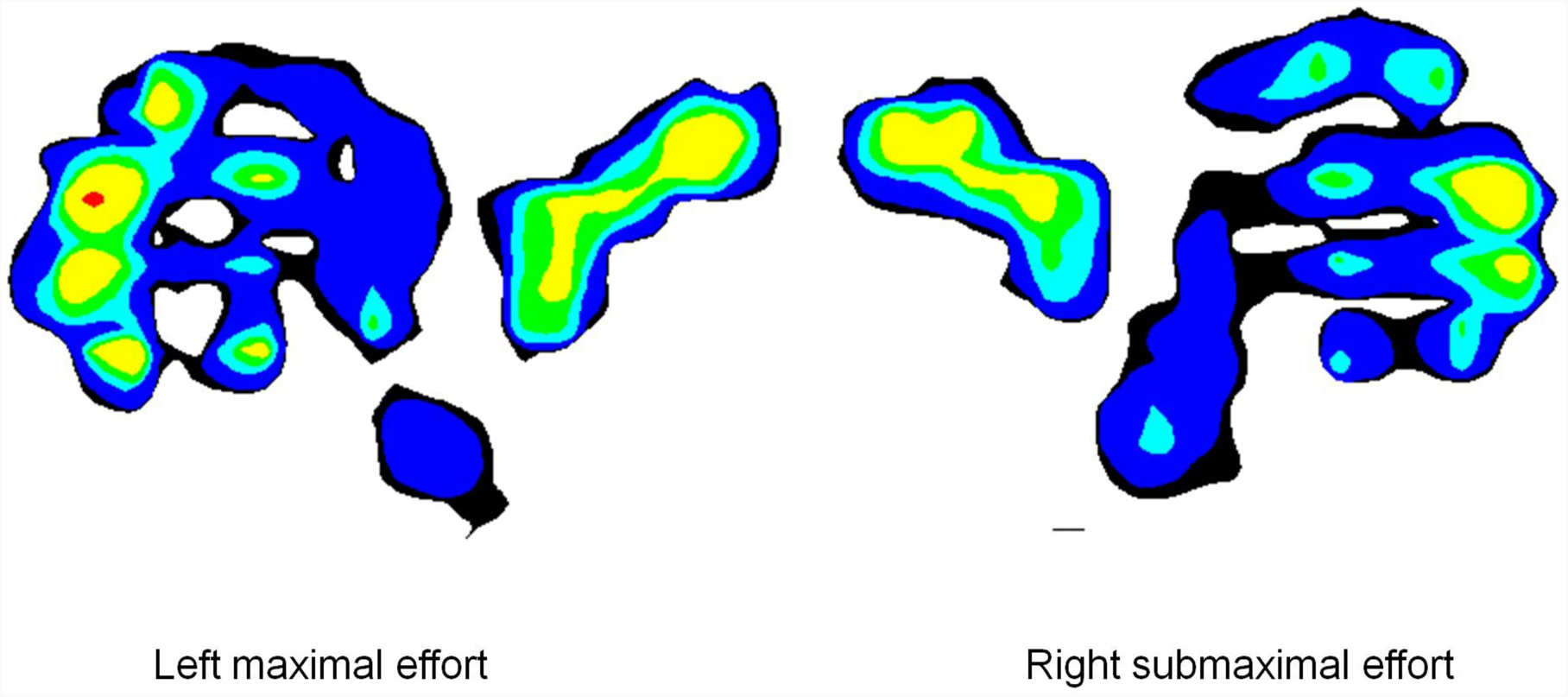
Grip pattern. Left side, maximal grip; right side, submaximal grip.

Principally, the results are pattern as shown in Fig 1. Analyzing such grip patterns using pattern recognition methods may be suitable for predicting feigned effort.

Machine learning (ML) is a field of computer science that evolved from artificial intelligence and pattern recognition. ML includes computer programs that are able to learn patterns by themselves and use them to make predictions. Nowadays, ML is widely used in, e.g., spam detection, word prediction on tablets, fraud detection in credit card transactions and weather prediction. In medical science, ML is used in genetic research, but is also rising in other fields, such as gait analysis (Christian et al. 2016). There are no studies on ML in grip pattern classification. Thus, the present study establishes machine learning as a tool for recognizing insincere effort in measuring grip strength.

## Methods

### Participants and data acquisition

As ML provides better results with a larger database, we used the data of several independent studies. All studies were supervised by the same researcher and used the same testing protocol. The grip force and load distribution were assessed using the novel (R) manugraphy system (novel biomechanics laboratory), Munich, Germany). The participants gripped a 20 cm circumference (64 mm diameter) cylinder while sitting on a stool without a backrest with the shoulder in neutral rotation, the arm adducted and the elbow flexed 90°. The cylinder was held in an upright position. The participants gripped the cylinder with maximum force for 5 sec. three times with 10 sec. breaks in between. Details of the manugraphy testing protocol are described here (Mühldorfer-Fodor et al. 2014).

Seventy-six healthy subjects performed grip force testing of both hands three times each on three independent testing days, which resulted in 1368 measurements.

In addition, 54 healthy subjects did grip force testing of both hands on two different days. These subjects were instructed to use one hand with maximum force and the opposite hand with submaximal effort to pretend weakness. The side that had to perform submaximal effort on the first session was randomized. Sides changed on the second visit. This protocol resulted in 324 measurements each of maximal and submaximal effort. In summary, a total of 2016 measurements could be used for further analysis, consisting of 1692 data records with maximal effort and 324 data records with submaximal effort (Fig. 2).

**Fig 2:**
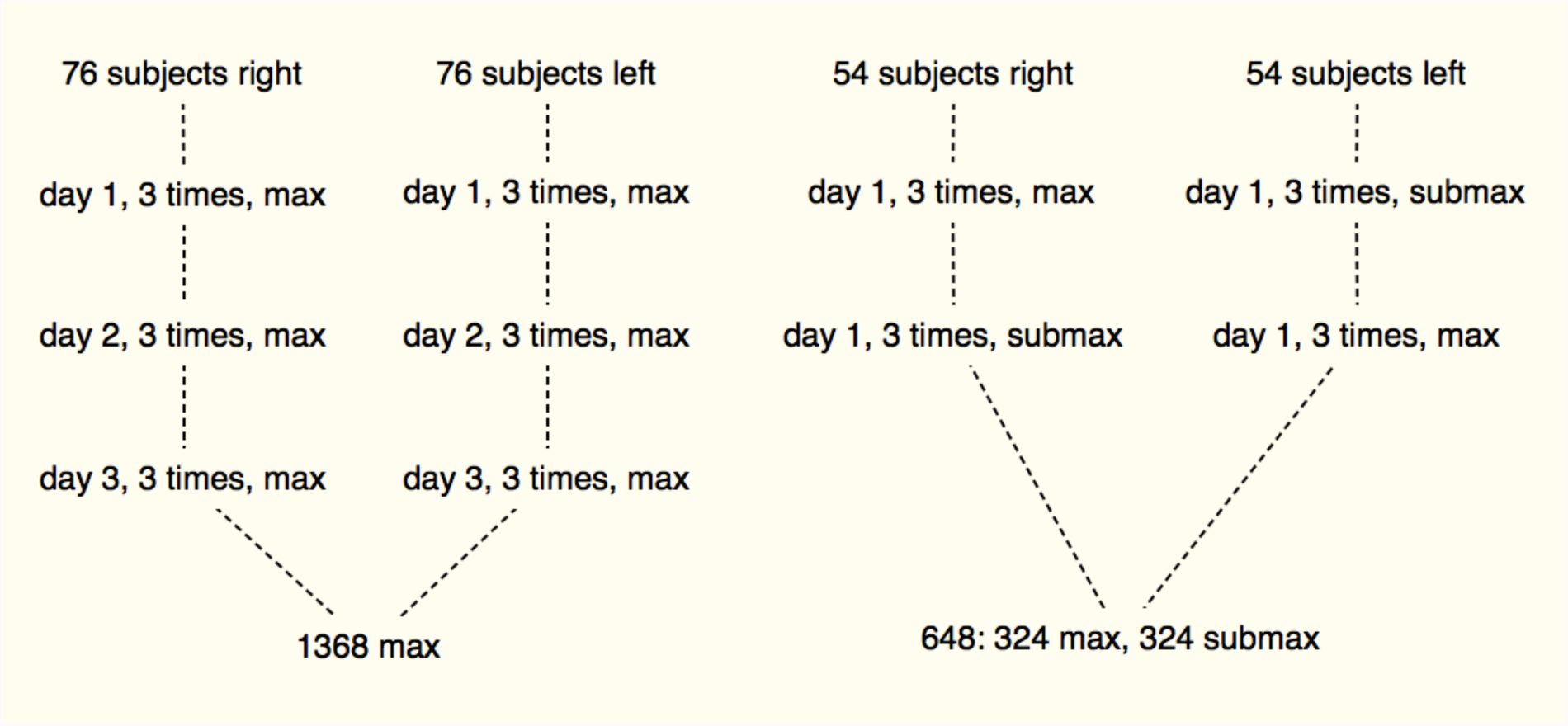
Flow diagram of measurements.

### Technical setting

The cylinder of the manugraphy system is covered with 896 capacitive sensors (28 x 32), with a spatial resolution of 2 sensors/cm2. Each sensor was individually calibrated to a maximum pressure of 600 kPa with an accuracy of < 5 %. The load distribution of the hand, not including the grip force values, was used for further analysis.

The average load applied to all the sensors in the sensor matrix was retrieved as a 2D raster diagram (Fig. 1). This shows the contact area of the hand gripping the cylinder in the form of a “load distribution map.” The contact area of the hand was sectioned by one investigator into 20 areas of interest (AOI) corresponding to the contact surfaces of each phalanx of the fingers and thumb (14), the Metacarpo-phalangeal joints (MCP) of the fingers (4), the thenar and the hypothenar region (Fig. 3). The force applied across the whole contact area was set at 100%, and the percentages of each of the 20 sections’ contributions to the total load were calculated.

**Fig 3:**
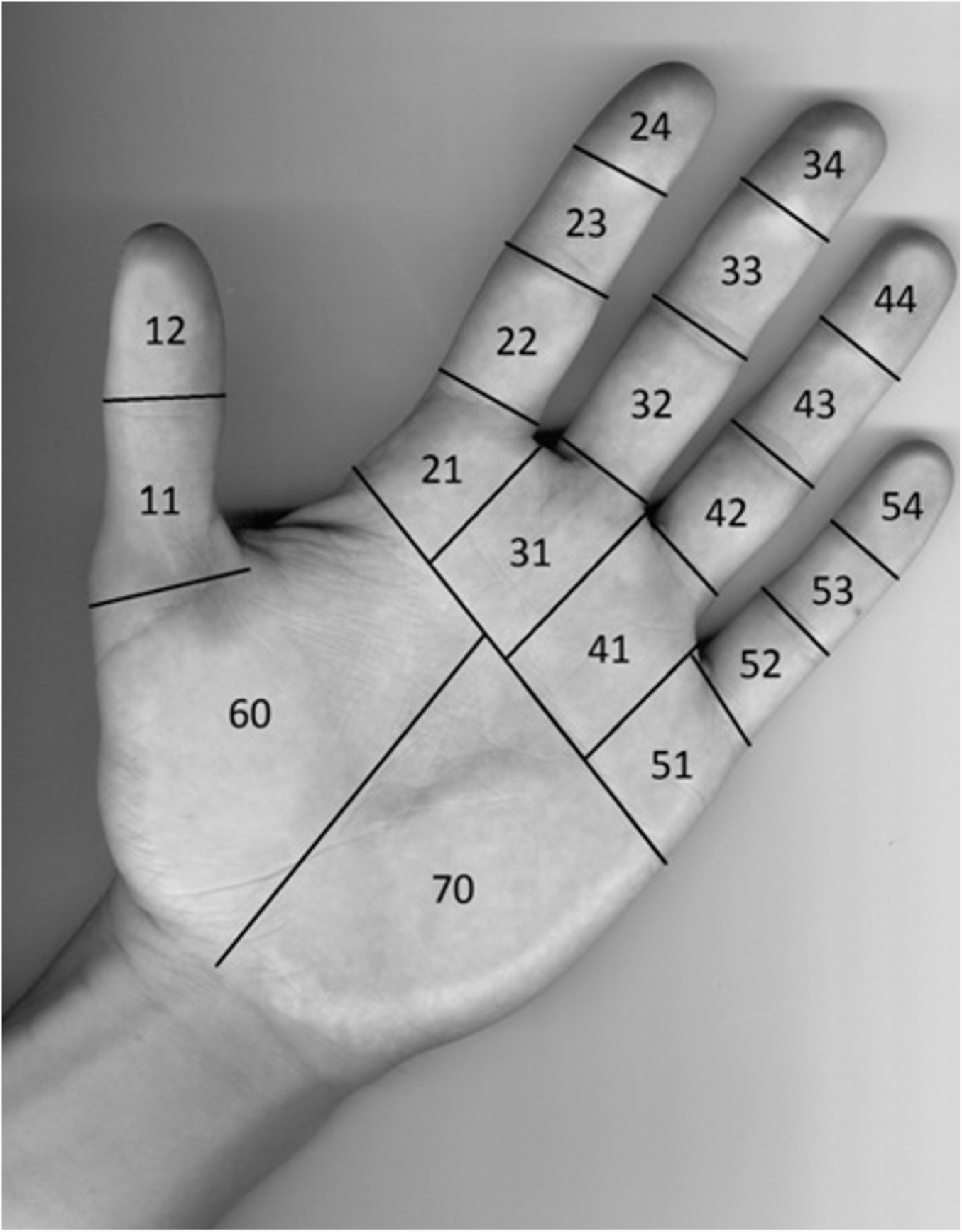
Representation of the 20 contact surfaces (Areas of Interest [AOI]) that are used for further analysis.

### Data analysis

#### Preprocessing

The data analysis was implemented in R (R Core Team 2013). All data were read from Excel files provided by the principal investigator of the different studies. Missing values were removed and categorical variables containing text were changed to numeric ones, as most ML programs can only deal with numeric input.

#### Statistical analysis

As stated above, there is a difference between maximal and submaximal grip. However, due to distribution of the data, submaximal effort cannot be predicted based on the statistical analysis (Fig. 4).

**Fig 4:**
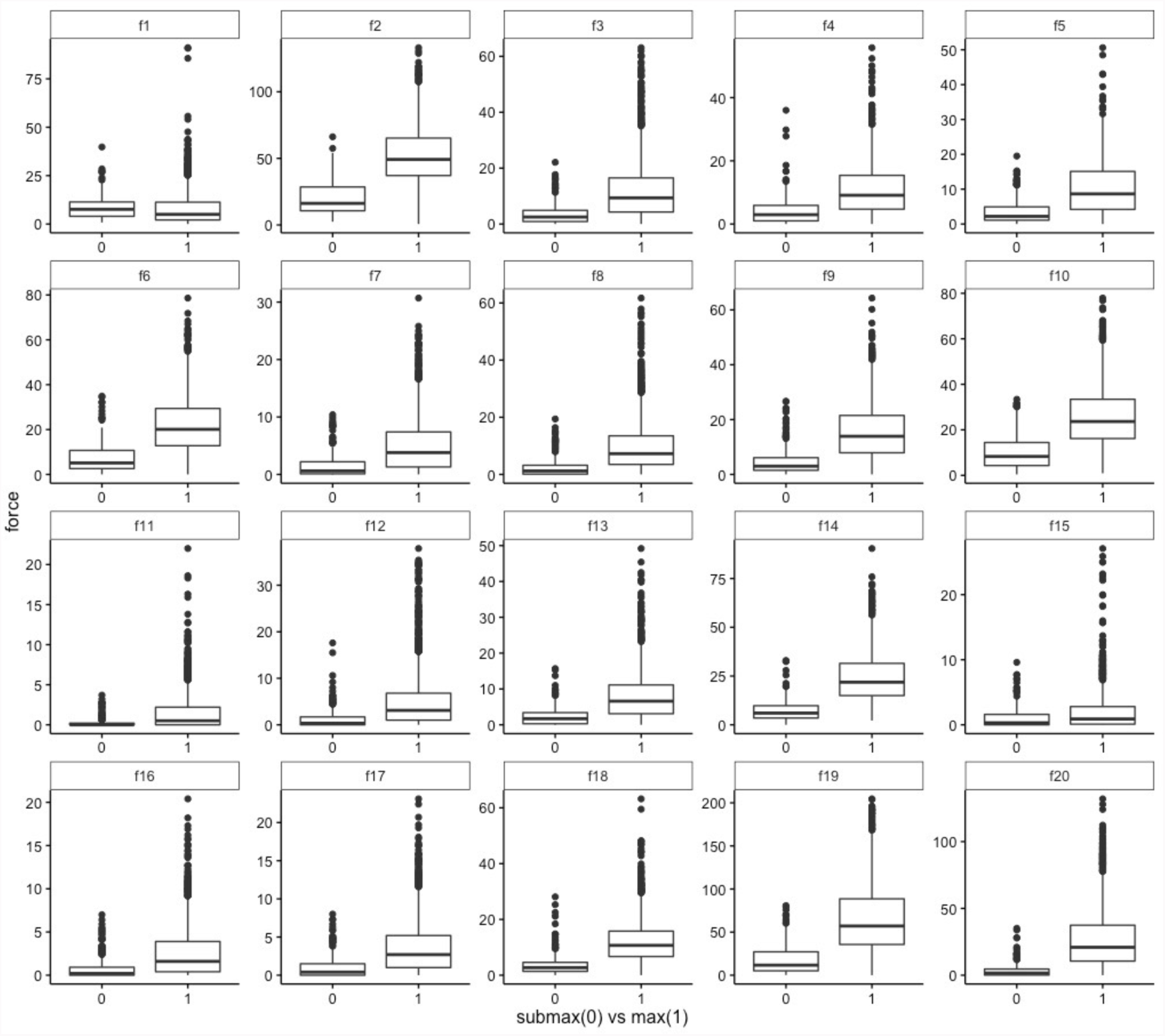
Boxplots (median, 25th and 75th percentile and outliers) of the individual areas of interest (AOI); x axis effort, y axis load

#### Feature extraction and building

The best results can be achieved by choosing good features to represent the data. This can often be achieved by having good knowledge of the underlying problem. We built seven fields of knowledge (FOK) representing the thumb, all fingers, the thenar and the hypothenar by summarizing the force of the respective AOI (Fig. 5). Due to skewness of the data, all FOK were converted to their logarithm. Additional features were built by combining existing features.

**Fig 5:**
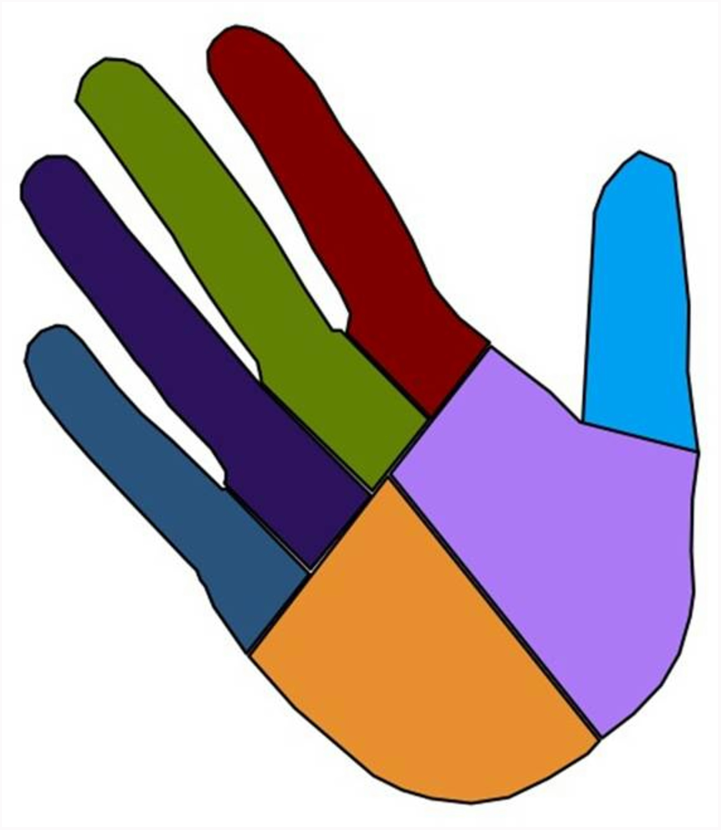
Graphical representation of fields of knowledge (FOK) calculated from the AIO.

Finally, a data table consisting of 2016 measurements with 12 features (Table 1) was used for training and testing the model.

**Tab 1:**
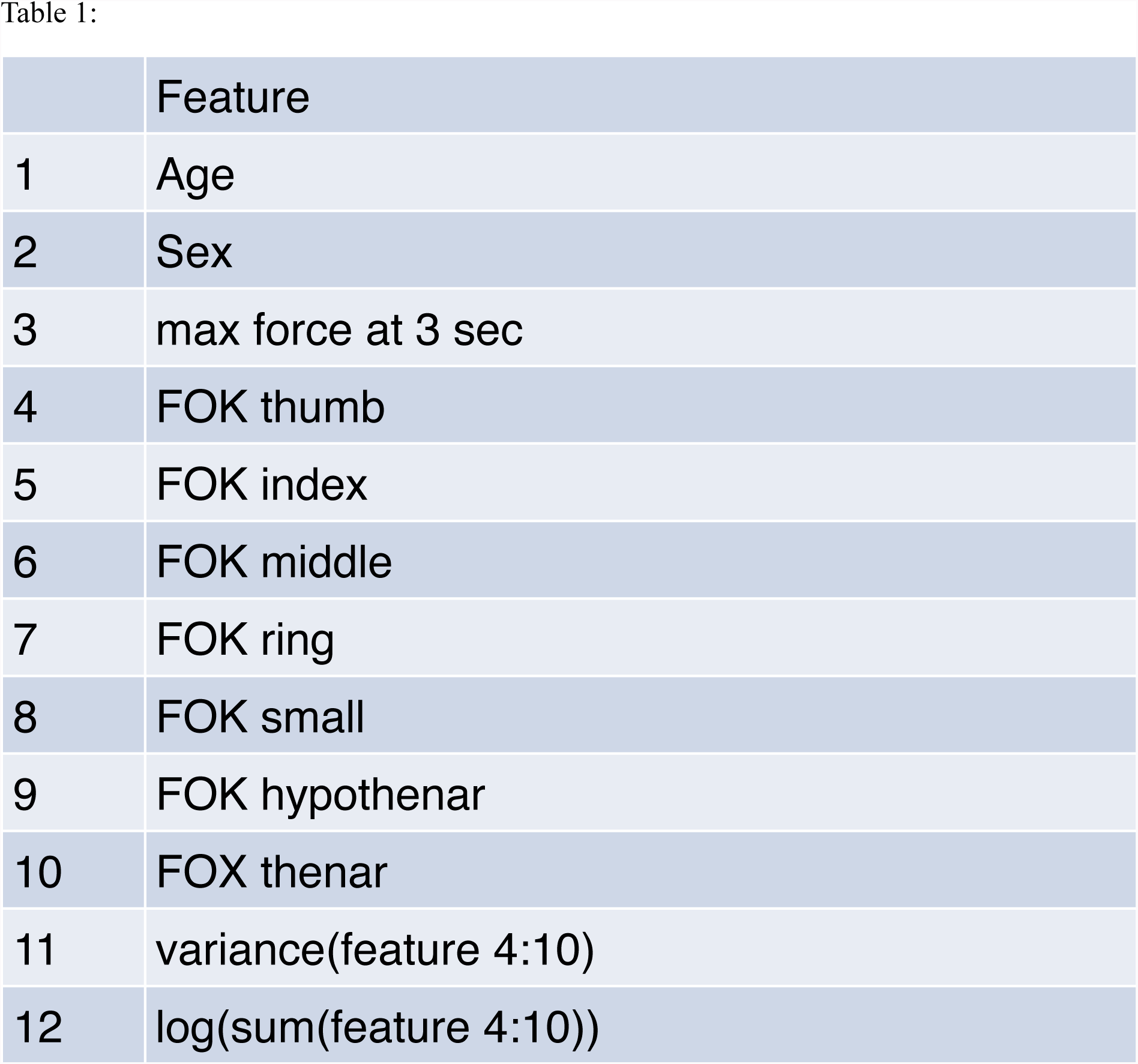
Features used for training. Features 4 to 10 are the fields of knowledge (FOK) explained in the text.

#### Model validation

Two subsets were built from the 2016 observations: 80% for training of the model and 20% for validation. Both data sets were chosen randomly. They were equally distributed in terms of maximal and submaximal observations.

During model training, cross validation was performed to avoid overfitting of the model. Finally, each model was tested against the validation data (Fig. 6). Before testing, identification of each effort was removed

#### Training

After testing several machine learning models and determining their performance based on the area under curve (AUC) from a receiver operation characteristic (ROC), it was found that the best results could be obtained using XGBoost. XGBoost is a tree-based model and a variant of the gradient boosting machine (GBM) (Chen & Guestrin 2016).

**Fig 6:**
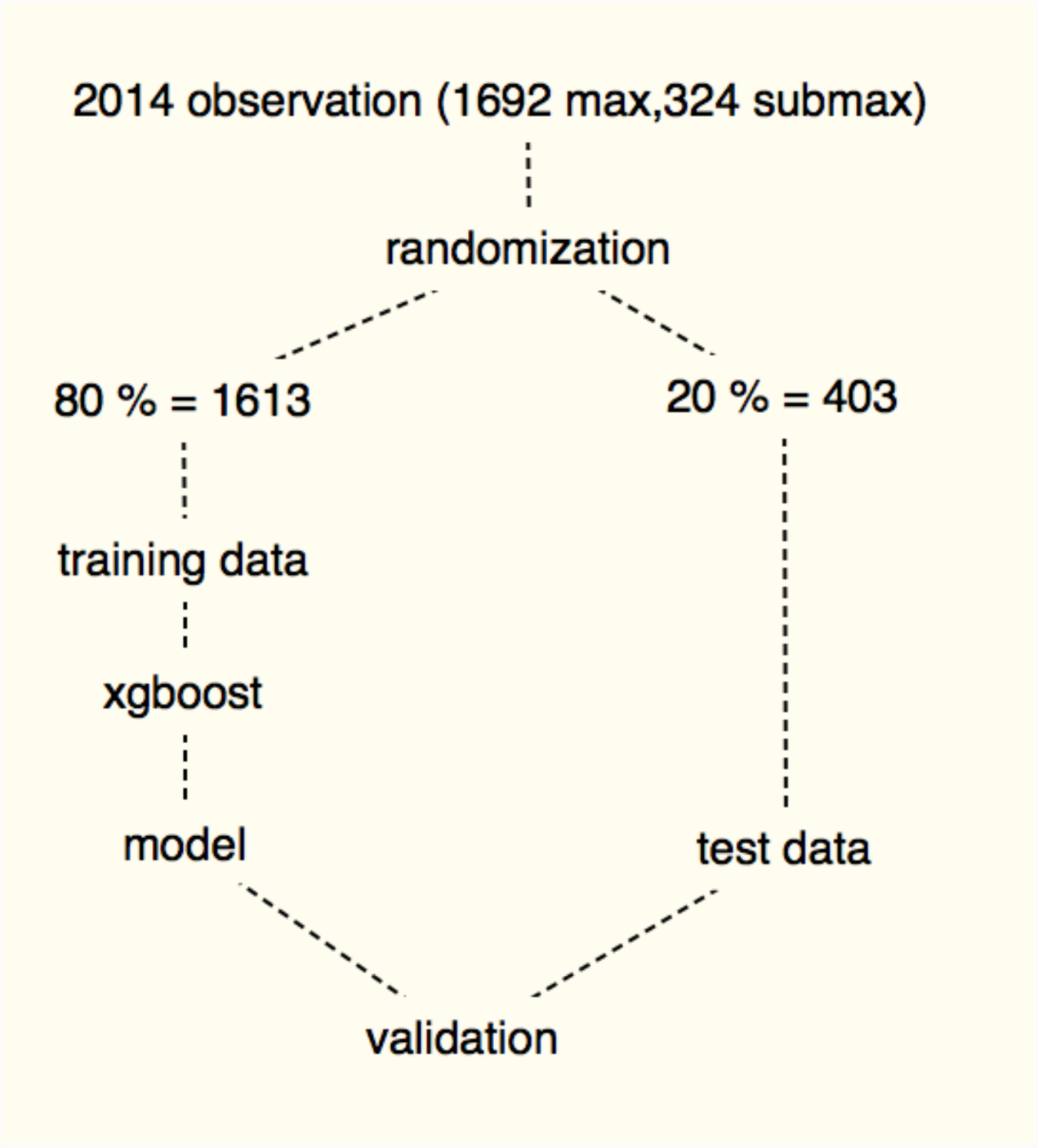
Evaluation of model validity

## Results

All results are based on the assumption that submaximal effort (0 =true positive) should be detected.

The results were calculated as probabilities rather than binary decisions. The ROC of the test data was plotted, and sensitivity and specificity at different thresholds were calculated from the data of the ROC (Fig. 7).

**Fig 7:**
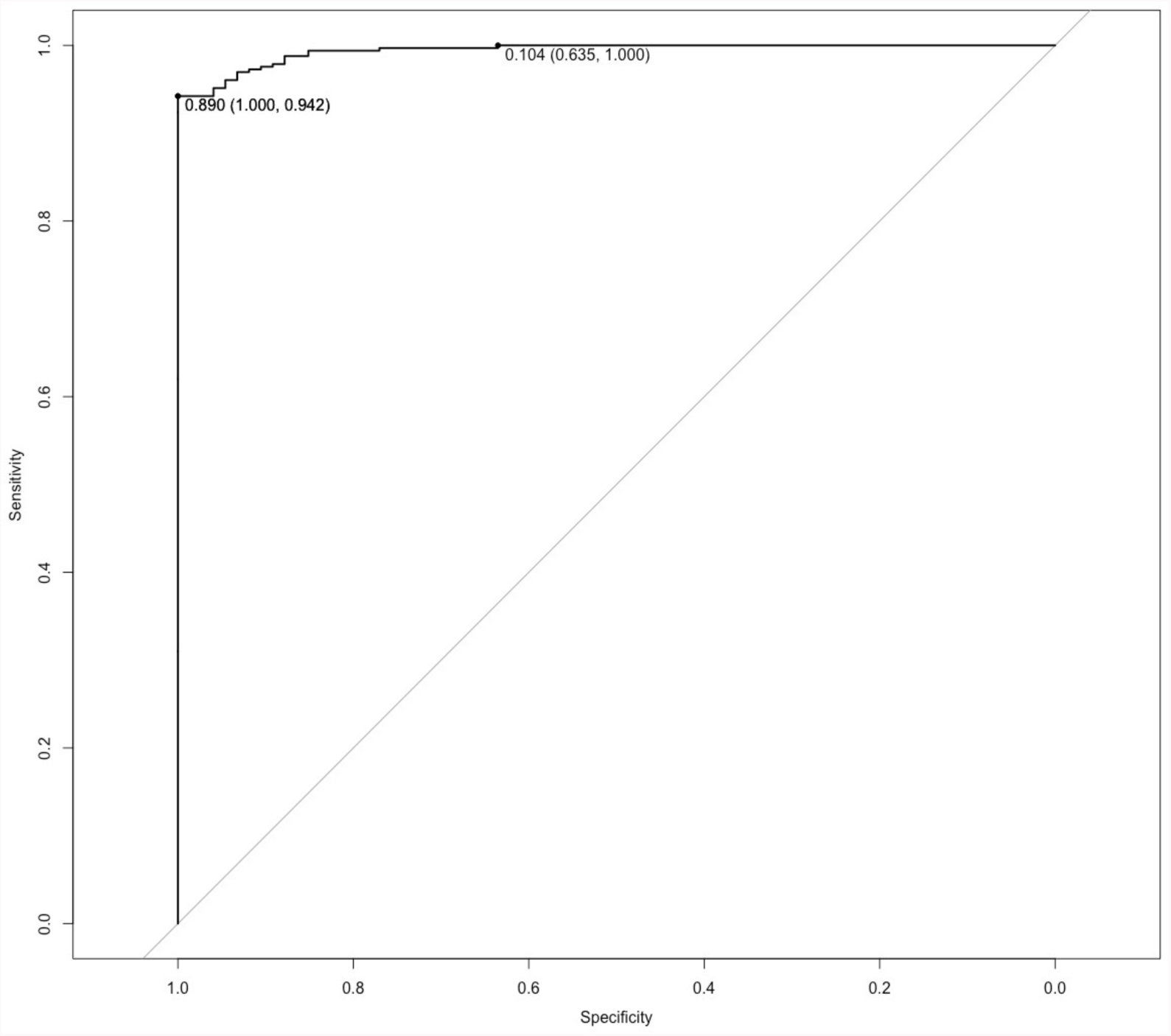
Receiver operation characteristic (ROC) for test data. Left point (specificity = 100%,sensitivity = 94.2 %, threshold: 0.89) and right point (sensitivity = 100%, specificity = 63.5%, threshold: 0.104)

Three points of the ROC were of special interest: the point that closest to the top-left corner of the curve (100% sensitivity and 100% specificity), the point with maximum specificity and the corresponding sensitivity, and the point with maximum sensitivity and the corresponding specificity. In our model, the first two points were the same.

The final model, which predicted submaximal effort, resulted in a sensitivity of 94.2% and a specificity of 100% with an accuracy of 95% (p < 0.0001) at a threshold of 0.890, corresponding to points one and two mentioned above. Assuming a prevalence of 20% insincere effort in patients, the PPV and NPV were calculated as PPV = 80% and NPV =100%.

Best sensitivity was calculated to be 100% at a specificity of 63.5%.

## Discussion

This study introduces a new approach for detecting submaximal effort in grip strength. The two classes, maximal and submaximal effort, can be predicted with high accuracy.

First, our results must be compared with the results of other studies. Dvir 1999 investigated isokinetic dynamometry and reported a sensitivity of 73.3% and a specificity of 97.05% at a confidence level of 95%. Shechtman, Sindhu, et al. 2007 evaluated the force-time curve of a special dynamometer and revealed a sensitivity and specificity of 0.93 vs 1.0 in males and 0.83 vs 0.93 in females. Tests using the JAMAR dynamometer as the five-rung test or the variation of measurements have low accuracy (Sindhu & Shechtman 2011; Shechtman 2001). Our study has the highest accuracy of all available studies in this field of knowledge.

Based on the threshold, we can have either better sensitivity or better specificity (Fig. 7).If we pair high sensitivity with low specificity, we might be able to identify all patients who made insincere efforts, but we would also accuse some patients who made sincere efforts of being insincere, which could result in serious problems such as decreased financial compensation or incorrect treatment. In our opinion, high specificity and lower sensitivity better fits the goals of healthcare. We will miss some patients who feigned effort, but we will not accuse patients who made sincere efforts of being insincere.

The value of a prediction model highly depends on the prevalence of the observed entity. If we assume a prevalence of 20% in a patient population, our prediction will miss approximately 20% of patients who made an insincere effort (PPV = 80%), but we will not falsely accuse anyone of feigning effort (100%). As mentioned above, this should be the viewpoint of physicians and therapists. The views and interpretations of insurance companies may be different.

Our study has several shortcomings. First of all, this method depends on load distribution assessment by the manugraphy system, which is not widely available. The measurements and data analysis are still time consuming. The analysis of the AOIs is difficult and can be biased if more investigators or less experienced investigators are involved. In a next step, we will simplify analysis of the manugraphy data. Finally, we want to analyze all of the raw data using ML.

For this study, the data only included healthy people. Further investigations are necessary that include patients with different pathologies of the hands, wrists or upper extremities.

## Conclusion

The present study showed the potential for machine learning to recognize grip patterns of sincere and insincere effort. The evaluation of the model revealed good performance on the test data. Even though there were some limits because only healthy adults were investigated, the large sample size enhanced the informative value. This approach could become a new method in clinical settings as well as scientific research. Further research is needed to refine data analysis and usefulness in patients with different pathologies.

